## Supplementary Figures for "Context-dependent effects explain divergent prognostic roles of Tregs in cancer"

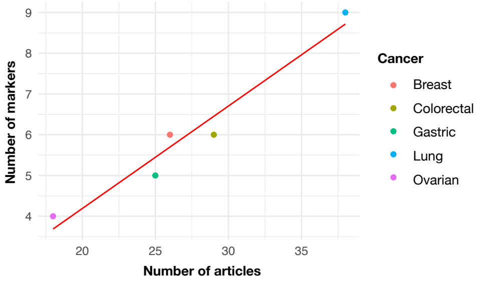


Supplementary Figure 1: Correlation between the number of markers used and the number of articles, by cancer type.

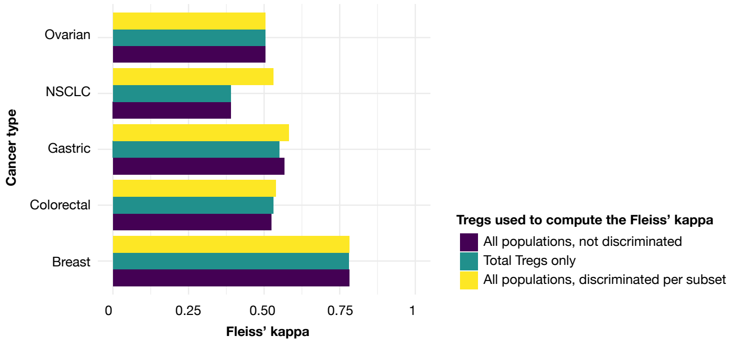


Supplementary Figure 2: Fleiss' kappa to determine the degree of agreement between articles, stratified by regulatory subset.

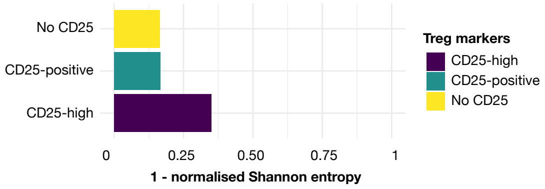


Supplementary Figure 3: Shannon entropy for articles defining Tregs based on expression of the CD25 marker.

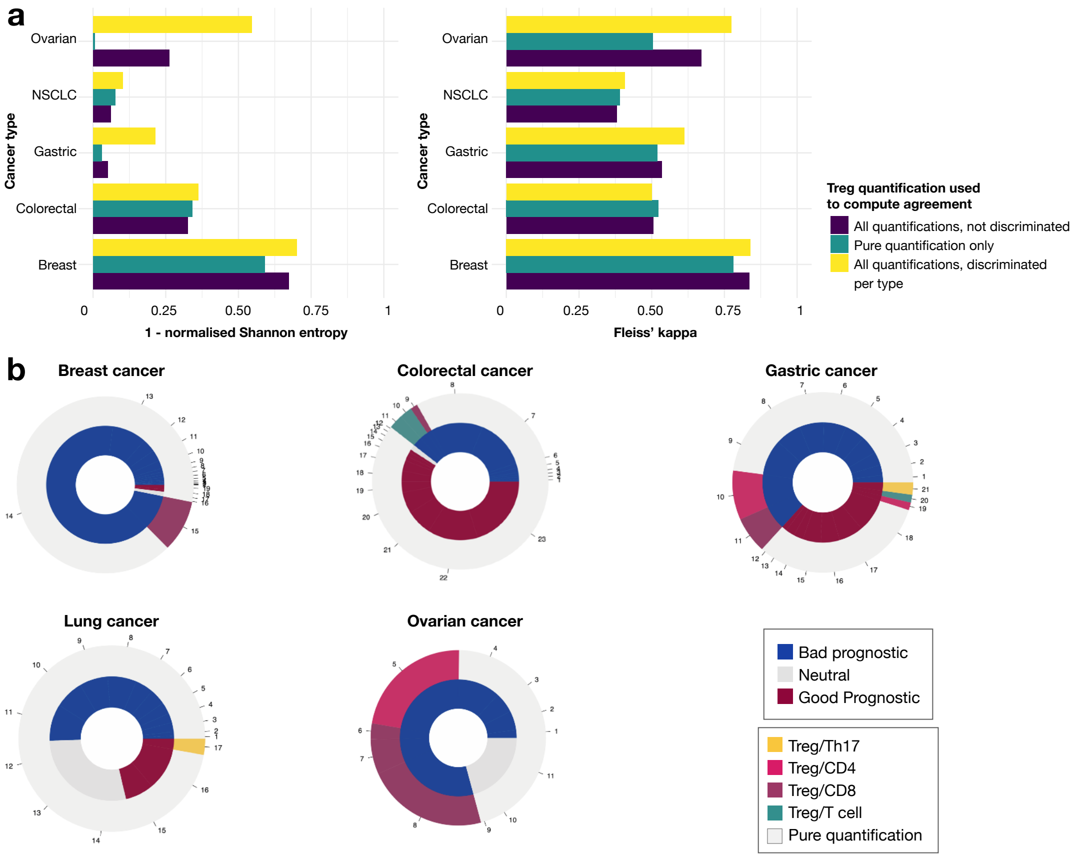


Supplementary Figure 4: Impact of the mode of Tregs quantification on the prognostic value. **(a)** Agreement between articles, stratified by type of quantification: Shannon entropy (left), Fleiss' kappa (right). **(b)** Pie chart of prognosis with information about quantification, by cancer type. Each portion is an article and its size reflects the number of patients included.

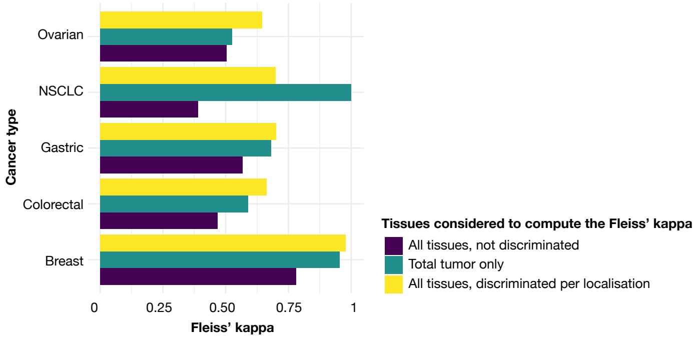


Supplementary Figure 5: Fleiss' kappa for assessing the degree of agreement between articles, stratified by anatomic location.

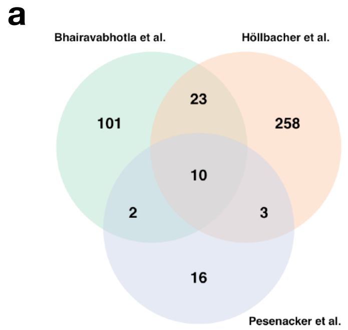


**b**

| Surface proteins | Intracellular proteins |
|---|---|
| **TNFRSF9**, CSF2RB, LRRC32, IL1R1, **CTLA4** | **FOXP3**, TRIB1, **IKZF2** (Helios), HPGD, STAM |

Supplementary Figure 6: Overlap between transcriptomic regulatory signatures. **(a)** Venn diagram of the gene signature for Tregs from.[46,47,45] **(b)** Genes at the intersection of the three signatures. Genes in bold have been repeatedly linked to the regulatory phenotype.

| Cancer | Good prognosis | Neutral prognosis | Poor prognosis | Undetermined |
|---|---|---|---|---|
| Breast | 28233108,<br>23075422 | 22842982,<br>25849846 | 24124553,<br>18820666,<br>17135638,<br>27851913,<br>23529839,<br>20181533,<br>23836289,<br>18413832,<br>23026134,<br>21717105,<br>22842982,<br>21521526,<br>27566250,<br>24562936 | 28388539,<br>17135638,<br>22760213,<br>27851913,<br>23529839,<br>20181533,<br>22836755,<br>19855964,<br>22116346,<br>23712790,<br>18294387 |
| Colorectal | 20386463,<br>19064967,<br>24005418,<br>19856313,<br>24675384,<br>24997850 | 23382847,<br>16740757 | 25268580,<br>26298011,<br>19064967,<br>21915633,<br>24005418,<br>22907255,<br>19577568,<br>19908042,<br>31681276 | 17205133,<br>22276195,<br>22319577,<br>23382847,<br>23613769,<br>25268580,<br>24064667,<br>20952660,<br>24005418,<br>22907255,<br>27851914,<br>22207629,<br>18985040,<br>25405854 |
| Gastric | 23807713,<br>24331841,<br>24170095,<br>21347781,<br>19732435,<br>32204925,<br>266799288 | | 29804142,<br>28817117,<br>27756099,<br>24657498,<br>24040244,<br>22374482,<br>22083420,<br>21792941,<br>20221835,<br>19900843,<br>19153062 | 29804142,<br>28817117,<br>26782287,<br>24261990,<br>24170095,<br>22083420,<br>21528082,<br>21347781,<br>20422211,<br>19900843,<br>19153062,<br>18224687,<br>18087278 |
| Lung | 22300751,<br>23335103,<br>279767333 | 22300751,<br>23305175,<br>23891508 | 17099880,<br>20234320,<br>21719142,<br>22363469,<br>22608141,<br>23305175,<br>23891508,<br>27773662,<br>27851914,<br>279767331,<br>279767332 | 15846066,<br>16698419<br>17163448,<br>17825949,<br>18771959,<br>19148592,<br>19332094,<br>19597336,<br>21258248,<br>21611754,<br>21663645,<br>22363469,<br>24345703,<br>24780112,<br>26042578,<br>26280204,<br>26541534,<br>27000869,<br>27474372,<br>27866241,<br>28513867,<br>28731226 |
| Ovarian | | 26077607,<br>20006900,<br>18314181 | 27748885,<br>26298430,<br>25365237,<br>24244610,<br>17875732,<br>16344461,<br>32902402 | 28437737,<br>27759594,<br>26482613,<br>25514665,<br>25416072,<br>23948613,<br>22865582,<br>22798340,<br>18166500,<br>18036640 |

Supplementary Table 1: List of articles used in this analysis with PMID references
